# A biomechanical investigation of stair descent in postmenopausal women: A regression analysis considering walking speed and T-score

**DOI:** 10.1101/2025.08.07.669220

**Authors:** Ali Dostan, Catherine A Dobson, Natalie Vanicek

**Affiliations:** Department of Engineering, Faculty of Science and Engineering, Nottingham Trent University; School of Engineering and Innovation, Open University; School of Sport, Exercise and Rehabilitation Sciences, Faculty of Health Sciences, University of Hull

**Keywords:** Stair descent biomechanics, Postmenopausal women, Bone mineral density (BMD), Osteoporosis, Eccentric muscle control, Gait analysis

## Abstract

**Introduction:** Previous studies have examined gait biomechanics during stair descent in older adults, but limited evidence exists on how bone mineral density (BMD) influences these mechanics in postmenopausal women, a group at increased risk of falls and fractures. This study investigated the relationship between self-selected stair descent speed and femoral neck BMD (T-score) with gait biomechanical parameters in postmenopausal women across a wide range of T-scores, from normal to osteoporotic.

**Methods:** Forty-five postmenopausal women (mean ± SD age: 67.3 ± 1.5 years) descended a five-step, custom-built staircase at a self-selected speed, without using handrails. Three-dimensional kinematics and ground reaction force (GRF) data were collected using motion capture and force plates. Multiple linear regression models assessed how stair descent speed and femoral neck T-score influenced stair descent gait parameters.

**Results:** The participants’ mean stair descent speed was 0.80 ± 0.21 m·s⁻¹. Speed significantly explained variance in temporal-spatial, kinematic, GRF, joint moment, and joint power parameters (R² = 7% to 49%, P ≤ 0.01). The inclusion of femoral neck T-score in the regression models further improved the explanatory power of the model in anterior pelvic tilt (R² = 21%, P ≤ 0.01), hip adduction (R² = 11%, P ≤ 0.01), hip extension (R² = 29%, P ≤ 0.01), knee flexion (R² = 21%, P ≤ 0.01), ankle dorsiflexion (R² = 19%, P ≤ 0.001), mid-stance vertical GRF (R² = 20%, P ≤ 0.01) during the forward continuance phase, and the second vertical GRF peak (R² = 15%, P ≤ 0.01).

**Conclusion:** This study demonstrates that stair descent speed and femoral neck T-score significantly influence lower limb biomechanics, accounting for up to 29% of the variance in key variables, particularly during the controlled lowering phase. Postmenopausal women with low BMD adopted altered movement strategies, such as increased frontal plane motion at the hip and pelvis, which may serve to maintain balance but could also elevate the risk of falls. The substantial eccentric muscle demands during stair descent contribute to mechanical loading of the lower limb skeleton, underscoring its potential not only as a complex functional task but also as an osteogenic stimulus. These findings highlight the value of targeted interventions aimed at improving eccentric strength and trunk control to enhance stair negotiation safety and support both musculoskeletal function and bone health in individuals with low BMD.

## Introduction

Stair descent imposes greater biomechanical demands than both level walking and stair ascent [1,2], primarily due to the need for controlled deceleration via eccentric muscle activity [3]. Falls on stairs can often lead to serious physical consequences, such as fractures (especially to the hip or wrist), head injuries, soft tissue damage, and long-term mobility issues [4] sometimes requiring hospitalisation [5]. This risk is increased in older adults, especially women, due to decreased musculoskeletal capacity, compounded by factors such as impaired balance, diminished sensory function, and delayed reaction time [6].

Postmenopausal women, particularly those with low bone mineral density (BMD), are at increased risk of developing osteoporosis later in life and are more susceptible to fractures [7]. Women over the age of 65 have a significantly higher risk of sustaining fractures in the hip, spine, and distal forearm compared to their younger peers [8]. According to the International Osteoporosis Foundation, the economic burden of osteoporosis in the UK is projected to reach £6 billion by 2030.

Understanding how women with low bone mineral density (BMD) or osteoporosis negotiate stairs will inform the development of targeted strategies to reduce falls risk [9]. Weight-bearing exercise is widely recommended as a key intervention to enhance physical capacity in older adults and to mitigate age-related bone loss in women [10]. Therefore, understanding the musculoskeletal contributions for safe stair walking is important when making evidence-based recommendations for appropriate exercises in this population [11].

Previous studies have explored age-related gait changes during stair descent in healthy older adults [2,12–14]. Older adults commonly demonstrate slower stair descent speeds, which have been associated with reduced knee flexor and extensor strength [15], and diminished ankle plantarflexor moments during early stance [2]. The biomechanical demands of stair descent often exceed the maximal isometric capacity of the hip and knee joints in older adults [16]. This is coupled with 26% greater stiffness in the lower limbs compared to younger adults possibly as a strategy to maintain the support leg more extended, because of reduced muscle strength, during stair descent [15]. These factors collectively raise concerns regarding the risk of falls and subsequent fractures during stair negotiation.

To compensate for functional losses, older adults adopt strategies such as redistributing joint moments proximally and utilising a higher proportion of muscle strength during stair descent [2]. Strategies, such as increasing the width of the base of support or redistributing joint moments, may enhance stability during stair descent [9,17]. These adaptations highlight the significant risks and challenges older adults encounter during stair descent, underscoring the need to understand and address these factors to prevent falls and fractures effectively.

A few studies have explored the effects of low BMD on the gait biomechanics of older women during level walking [18–25]. Some of these studies only examined the temporal-spatial parameters [21,22]. Moisio et al (2004) reported that 20% of the variance in femoral neck BMD was explained by the internal hip joint moment. However, when body mass was taken into consideration [19], no significant relationship was found between the two variables. Another study identified hip adductor and extensor moments, along with several peak joint powers, as significant predictors of low BMD in postmenopausal women [23]. However, ElDeeb and Khodair (2014) did not examine gait speed as a predictor, so it remains unclear how much gait speed alone contributed to variations in BMD among postmenopausal women. The influence of speed on gait variables during stair descent has been well established prior to this study [15,26]. There remains a lack of research examining the influence of gait speed and BMD (quantified by T-score) on the biomechanics of older women, particularly those with osteopenia or osteoporosis.

The aim of this study was to investigate the relationships between gait parameters during stair descent and key predictor variables, including stair descent speed and T-score, in postmenopausal women with a broad range of bone mineral densities. Understanding these relationships is essential for identifying musculoskeletal strategies during functional daily activities, such as stair descent, across the spectrum of bone health in older women. It was hypothesised that stair descent speed would account for the majority of variance in gait parameters, and that including T-score in the regression model would explain additional variance, particularly in sagittal and frontal plane hip kinematics and kinetics, as well as ankle biomechanics.

## Method

### Participants

Patient records of over 5,000 women from the local Centre for Metabolic Bone Disease were screened based on predefined eligibility criteria, resulting in the identification of more than 200 eligible participants. The likelihood of pathology-related changes was minimised through the application of strict eligibility requirements. Inclusion criteria stipulated participants were women aged 65-70 years, with a BMI of 18-30 kg/m^2^, who must have had a DXA scan within the previous 12 months, and with a femoral neck T-score of 0 to −4. Bone mineral density (BMD) was assessed using T-scores and classified into three categories: normal (T-score between −1.0 and +1.0 standard deviations [SD] of the young adult reference mean), osteopenia (T-score between −1.0 and −2.5 SD), and osteoporosis (T-score ≤ −2.5 SD).

All participants were required to be free of cardiac conditions and capable of ascending and descending a five-step staircase using a reciprocal gait pattern (without the use of handrails). Exclusion criteria included any diagnosed neurological disorders, observable gait abnormalities, or a history of treatment with glucocorticoids, teriparatide, bisphosphonates, or hormone replacement therapy within five years prior to data collection. Ethical approval for the study was granted by the Research Ethics Committee (reference: 711/YH/0347). Participants were fully informed about the study and gave their written consent before taking part. Participant demographics are summarised in Table 1.

**Table 1.** Demographic and Baseline Data of Participants (n = 45)

|  | Mean | SD | Range |
| --- | --- | --- | --- |
| Age (years) | 67.3 | 1.4 | 65 to 70 |
| Height (cm) | 161.4 | 4.9 | 151 to 172.5 |
| Mass (Kg) | 63.5 | 8.6 | 47.8 to 80.4 |
| BMI (Kg/m<sup>2</sup>) | 24.1 | 2.8 | 18.6 to 29.2 |
| Femoral neck T-score | -1.5 | 0.8 | 0.9 to -3 |
| Self-reported number of active days per week | 5 | 2 | 0 to 7 |
| Age of menopause (years) | 50 | 4.5 | 38 to 58 |
| Number of falls (last 12 months) | 1 | 1 | 0 to 3 |
| Number of fractures (>50 years) | 1 | 1 | 0 to 4 |

### Protocol

Three-dimensional lower limb kinematics during stair descent were captured using twelve high-speed Pro-Reflex MCU1000 infrared cameras (Qualisys, Gothenburg, Sweden) operating at a sampling frequency of 100 Hz. The cameras were positioned around a custom-built, five-step wooden staircase (step height: 20 cm; tread depth: 30 cm; width: 80 cm) equipped with two embedded force platforms (Model 9286AA, Kistler, Winterthur, Switzerland) located on steps 2 and 3 (**Error! Reference source not found.**) that captured ground reaction force (GRF) data at 500 Hz. Kinematic and kinetic data were synchronised via Qualisys Track Manager software (version 2.9, Qualisys, Sweden). The uppermost step of the staircase served as a landing platform, bordered by handrails to ensure participant safety and allow unrestricted turning. Participants wore tight-fitting shorts, a t-shirt, and flat, flexible footwear to minimise movement artefacts and maintain natural gait patterns. Retroreflective markers were affixed to anatomical landmarks on the pelvis and lower extremities following a six degrees of freedom (6DOF) marker configuration to enable precise segmental motion tracking [27]. Participants were instructed to begin from the rear edge of the landing platform and take one to two initiation steps before descending the staircase at a self-selected, comfortable pace. They were asked to avoid using the handrails unless required for safety. Following stair descent, participants continued ambulating for an additional 5 metres along a level walkway to ensure steady-state gait for data capture.

### Data analysis

Gait data were subsequently analysed in Visual 3D™ (version 3.0, C-Motion, Rockville, MD, USA) and normalised to the gait cycle starting with toe off on the top step. The gait sub-phases were labelled according to McFadyen and Winter (1988) as: leg pull-through and foot placement in swing; and weight acceptance, forward continuance and controlled lowering in stance [28]. Marker trajectory data were first gap-filled using cubic spline interpolation for minor occlusions and then low-pass filtered with a fourth-order, zero-lag Butterworth filter at a 6 Hz cut-off frequency, based on residual analysis.

Filtered marker trajectories were used to compute joint kinematics. Joint moments were normalised to body mass, and net joint moments were calculated using inverse dynamics [28]. Kinetic data, including GRFs and joint moments, were low-pass filtered with a 25 Hz cut-off frequency to effectively reduce high-frequency noise while preserving essential biomechanical signals, consistent with established protocols. Gait events were identified automatically in Visual3D and verified visually [29]. Data from the right and left limbs were averaged to represent overall lower limb function, as participants were healthy and no significant asymmetry was anticipated. Joint power bursts at the hip (H1–H3), knee (K1–K5), and ankle (A1–A3) were labelled according to McFadyen and Winter (1998).

### Statistical analysis

A sensitivity analysis was conducted using G*Power to determine the minimum detectable effect size for a multiple regression analysis. With a total sample size of 45 participants, two predictor variables, an alpha level of 0.05, and a desired power of 0.80, the analysis indicated that the minimum detectable effect size was f² = 0.18, representing a medium effect size according to Cohen’s (1988) guidelines. This suggests that the study was sufficiently powered to detect moderate or larger effects.

A histogram of residuals, together with assessments of skewness and kurtosis, was used to evaluate the normality of the data distribution. Multicollinearity among the three predictor variables was assessed using the variance inflation factor (VIF), with an average VIF of 1.07, indicating no evidence of problematic multicollinearity [30].

Statistical analyses were performed using Stata software (version 14, StataCorp, Texas, USA). Multiple linear regression analyses were conducted to examine the extent to which stair descent gait speed and femoral neck T-score explained variance (R²) in gait variables, including temporal-spatial parameters, joint kinematics, and kinetics. Stair descent gait speed was included as a predictor variable based on its well-documented association with gait parameters [24,31,32]. Similarly, T-score was included because it is frequently used in gait-related research to distinguish older women with osteopenia or osteoporosis from those with normal bone mineral density [18,19,21,23]. The regression models were developed using a blockwise-entry method. The first model included gait speed alone; the second included both gait speed and T-score. The slope coefficient (B) was reported to quantify the relationship between predictors and outcomes. Statistical significance was set at (P ≤ 0.05).

## Results

The mean (SD) stair descent speed was 0.80 (0.21) m·s⁻¹ (see Table 2). When stair descent speed was entered as the sole predictor, it explained a modest proportion of the variance (R² = 11–36%, P ≤ 0.05) in the temporal-spatial and kinematic parameters (Table 2).

**Table 2.** Temporal-spatial gait parameters during stair descent.

|  | Mean | SD | Range |
| --- | --- | --- | --- |
| Speed (m/s) | 0.81 | 0.17 | 0.56 to 1.24 |
| Stance time (% of GC) | 59 | 9 | 44 to 68 |
| Swing time (% of GC) | 42 | 4 | 32 to 56 |
| Cadence (steps/min) | 112 | 20 | 80 to 174 |

### Joint kinematics

The addition of T-score as a predictor in the second regression model significantly improved the explanatory power of the model for selected kinematic variables (P ≤ 0.01). Specifically, T-score accounted for increased variance in anterior pelvic tilt (R² = 21%) and hip extension (R² = 29%) during controlled lowering, hip adduction during leg pull-through (R² = 11%), knee flexion during controlled lowering (R² = 21%), ankle dorsiflexion during controlled lowering (R² = 19%), and overall ankle range of motion (R² = 18%) (Table 3, Figure 2).

**Figure 1.**
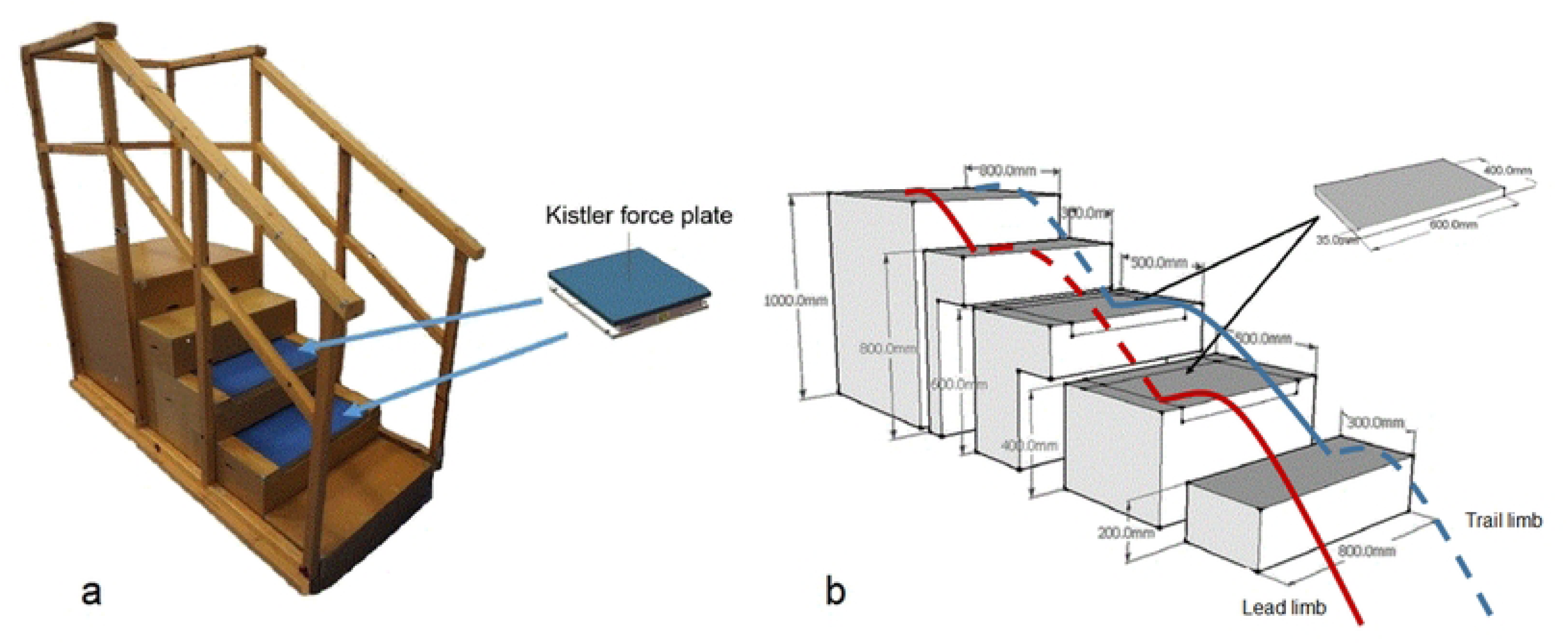
(a) Laboratory staircase with force plates embedded in the second and third steps (force plate 1 and force plate 2, respectively). (b) Schematic illustration of lead (red line) and trail (blue line) limb gait cycles during stair descent.

**Figure 2.**
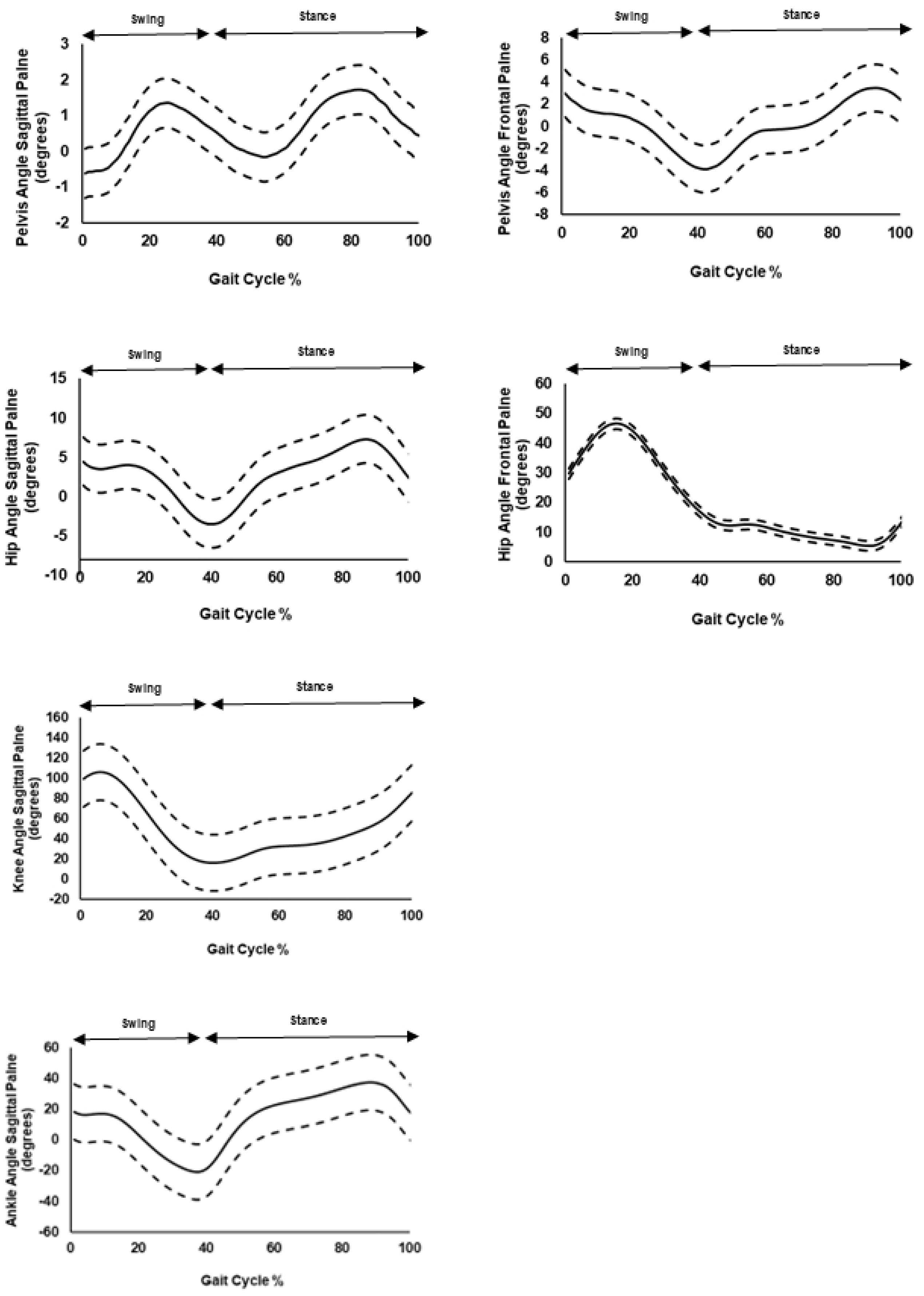
Ensemble averages (solid lines) ±1 standard deviation (dashed lines) of lower limb joint kinematics profiles during stair descent. Positive values represent pelvic hike, anterior pelvic tilt, hip adduction, hip and knee flexion, and ankle dorsiflexion.

**Table 3.**
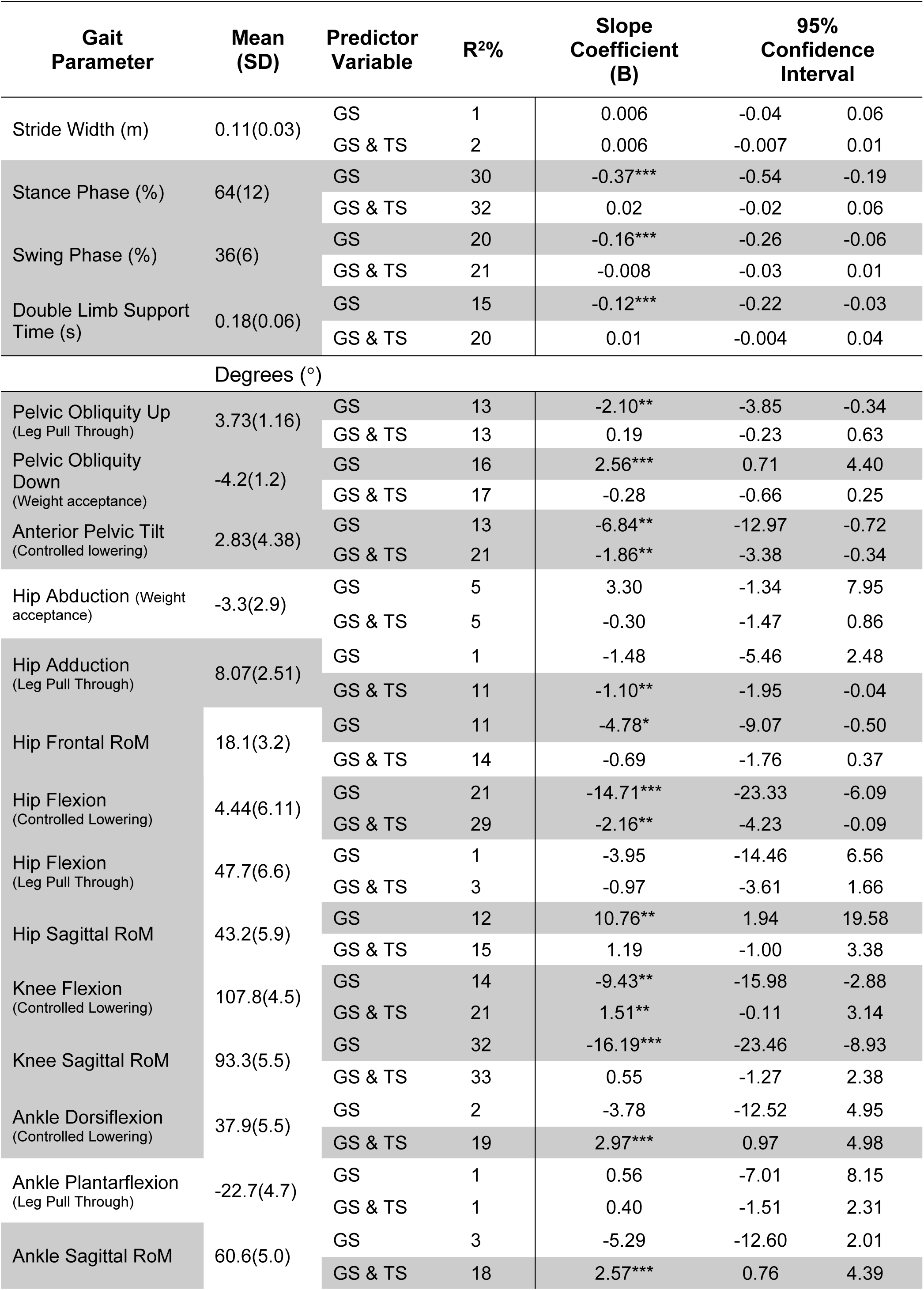
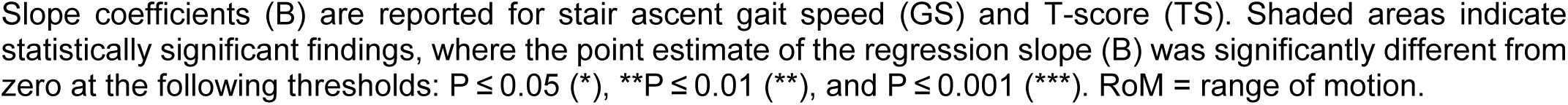
Explained variance (R²) and slope coefficients for temporal-spatial parameters and joint kinematics observed during stair descent.

| Gait Parameter | Mean (SD) | Predictor Variable | R<sup>2</sup>% | Slope Coefficient (B) | 95% Confidence Interval |  |
| --- | --- | --- | --- | --- | --- | --- |
| Stride Width (m) | 0.11(0.03) | GS | 1 | 0.006 | -0.04 | 0.06 |
|  |  | GS & TS | 2 | 0.006 | -0.007 | 0.01 |
| Stance Phase (%) | 64(12) | GS | 30 | -0.37*** | -0.54 | -0.19 |
|  |  | GS & TS | 32 | 0.02 | -0.02 | 0.06 |
| Swing Phase (%) | 36(6) | GS | 20 | -0.16*** | -0.26 | -0.06 |
|  |  | GS & TS | 21 | -0.008 | -0.03 | 0.01 |
| Double Limb Support Time (s) | 0.18(0.06) | GS | 15 | -0.12*** | -0.22 | -0.03 |
|  |  | GS & TS | 20 | 0.01 | -0.004 | 0.04 |
| Degrees (°) |  |  |  |  |  |  |
| Pelvic Obliquity Up (Leg Pull Through) | 3.73(1.16) | GS | 13 | -2.10** | -3.85 | -0.34 |
|  |  | GS & TS | 13 | 0.19 | -0.23 | 0.63 |
| Pelvic Obliquity Down (Weight acceptance) | -4.2(1.2) | GS | 16 | 2.56*** | 0.71 | 4.40 |
|  |  | GS & TS | 17 | -0.28 | -0.66 | 0.25 |
| Anterior Pelvic Tilt (Controlled lowering) | 2.83(4.38) | GS | 13 | -6.84** | -12.97 | -0.72 |
|  |  | GS & TS | 21 | -1.86** | -3.38 | -0.34 |
| Hip Abduction (Weight acceptance) | -3.3(2.9) | GS | 5 | 3.30 | -1.34 | 7.95 |
|  |  | GS & TS | 5 | -0.30 | -1.47 | 0.86 |
| Hip Adduction (Leg Pull Through) | 8.07(2.51) | GS | 1 | -1.48 | -5.46 | 2.48 |
|  |  | GS & TS | 11 | -1.10** | -1.95 | -0.04 |
| Hip Frontal RoM | 18.1(3.2) | GS | 11 | -4.78* | -9.07 | -0.50 |
|  |  | GS & TS | 14 | -0.69 | -1.76 | 0.37 |
| Hip Flexion (Controlled Lowering) | 4.44(6.11) | GS | 21 | -14.71*** | -23.33 | -6.09 |
|  |  | GS & TS | 29 | -2.16** | -4.23 | -0.09 |
| Hip Flexion (Leg Pull Through) | 47.7(6.6) | GS | 1 | -3.95 | -14.46 | 6.56 |
|  |  | GS & TS | 3 | -0.97 | -3.61 | 1.66 |
| Hip Sagittal RoM | 43.2(5.9) | GS | 12 | 10.76** | 1.94 | 19.58 |
|  |  | GS & TS | 15 | 1.19 | -1.00 | 3.38 |
| Knee Flexion (Controlled Lowering) | 107.8(4.5) | GS | 14 | -9.43** | -15.98 | -2.88 |
|  |  | GS & TS | 21 | 1.51** | -0.11 | 3.14 |
| Knee Sagittal RoM | 93.3(5.5) | GS | 32 | -16.19*** | -23.46 | -8.93 |
|  |  | GS & TS | 33 | 0.55 | -1.27 | 2.38 |
| Ankle Dorsiflexion (Controlled Lowering) | 37.9(5.5) | GS | 2 | -3.78 | -12.52 | 4.95 |
|  |  | GS & TS | 19 | 2.97*** | 0.97 | 4.98 |
| Ankle Plantarflexion (Leg Pull Through) | -22.7(4.7) | GS | 1 | 0.56 | -7.01 | 8.15 |
|  |  | GS & TS | 1 | 0.40 | -1.51 | 2.31 |
| Ankle Sagittal RoM | 60.6(5.0) | GS | 3 | -5.29 | -12.60 | 2.01 |
|  |  | GS & TS | 18 | 2.57*** | 0.76 | 4.39 |

Slope coefficients (B) are reported for stair ascent gait speed (GS) and T-score (TS). Shaded areas indicate statistically significant findings, where the point estimate of the regression slope (B) was significantly different from zero at the following thresholds: $P \leq 0.05$ (\*), $**P \leq 0.01$ (\*\*), and $P \leq 0.001$ (\*\*\*). RoM = range of motion.

### Ground reaction forces

Stair descent speed alone explained a significant portion of the variance in key vertical GRF parameters, including the mid-stance GRF (i.e., GRF trough) during the forward continuance phase (R² = 14%, P ≤ 0.01), the second GRF peak during the controlled lowering phase (R² = 7%, P ≤ 0.05), and the GRF decay rate following initial contact (R² = 7%, P ≤ 0.05) (Table 4). When femoral neck T-score was added to speed in the second regression model, the combined explanatory power significantly increased for the GRF in mid-stance (R² = 20%) and the second GRF peak (R² = 15%) (P ≤ 0.01 for both). These findings suggest that both gait speed and bone density status (T-score) contribute meaningfully to the mechanical loading experienced during specific phases of stair descent.

**Table 4.** Explained variance (R²) and slope coefficients (B) for peak ground reaction forces and joint moments during stair descent.

| Gait Parameter | Mean (SD) | Predictor Variable | $R^2\%$ | Slope Coefficient (B) | 95% Confidence Interval | |
| --- | --- | --- | --- | --- | --- | --- |
|  | (N.kg<sup>-1</sup>) |  |  |  |  |  |
| Peak posterior force<br>(Weight acceptance) | 0.01(0.02) | GS | 1 | -0.001 | -0.04 | 0.04 |
|  |  | GS & TS | 4 | -0.006 | -0.01 | 0.003 |
| Peak anterior force<br>(Controlled Lowering) | -0.05(0.05) | GS | 3 | -0.03 | -0.12 | 0.04 |
|  |  | GS & TS | 3 | -0.001 | -0.01 | 0.01 |
| 1<sup>st</sup> Vertical GRF peak<br>(Weight acceptance) | 1.55(0.18) | GS | 2 | 0.14 | -0.17 | 0.47 |
|  |  | GS & TS | 3 | -0.02 | -0.08 | 0.04 |
| Vertical GRF mid-stance<br>(Forward Continuance) | 0.75(0.11) | GS | 14 | 0.23** | 0.05 | 0.40 |
|  |  | GS & TS | 20 | -0.03** | -0.06 | 0.004 |
| 2<sup>ND</sup> Vertical GRF peak<br>(Controlled Lowering) | 0.95(0.12) | GS | 7 | 0.18* | -0.01 | 0.38 |
|  |  | GS & TS | 15 | -0.03** | -0.08 | 0.002 |
| Loading rate (N.kg<sup>-1</sup>.s<sup>-1</sup>) | 13.35(3.25) | GS | 4 | 4.21 | -1.61 | 10.04 |
|  |  | GS & TS | 8 | -0.74 | -1.96 | 0.42 |
| Decay rate (N.kg<sup>-1</sup>.s<sup>-1</sup>) | 5.8(1.7) | GS | 7 | 2.57* | -0.35 | 5.49 |
|  |  | GS & TS | 10 | -0.32 | -0.94 | 0.30 |
|  | (N.m.kg<sup>-1</sup>) |  |  |  |  |  |
| Hip Abductor<br>(Weight Acceptance) | 0.71(0.36) | GS | 13 | 0.65** | 0.12 | 1.19 |
|  |  | GS & TS | 13 | 0.004 | -0.12 | 0.13 |
| Hip Adductor<br>(Controlled Lowering - Leg Pull Through) | -0.1(0.04) | GS | 2 | -0.02 | -0.08 | 0.03 |
|  |  | GS & TS | 3 | -0.005 | -0.02 | 0.009 |
| Hip Flexor<br>(Leg Pull Through) | 0.63(0.3) | GS | 15 | -0.78*** | -1.35 | -0.20 |
|  |  | GS & TS | 17 | 0.02 | -0.06 | 0.11 |
| Hip Flexor<br>(Controlled Lowering) | -0.84(0.40) | GS | 7 | 0.64* | -0.10 | 1.38 |
|  |  | GS & TS | 7 | -0.05 | -0.14 | 0.03 |
| Knee Extensor<br>(Weight Acceptance) | 0.82(0.30) | GS | 15 | 0.57** | 0.10 | 1.04 |
|  |  | GS & TS | 16 | -0.03 | -0.13 | 0.06 |
| Knee Extensor<br>(Controlled Lowering) | 1.34(0.40) | GS | 6 | -0.50 | -1.12 | 0.12 |
|  |  | GS & TS | 7 | -0.05 | -0.20 | 0.09 |
| Knee Flexor<br>(Controlled Lowering) | 0.5(0.2) | GS | 2 | 0.15 | -0.22 | 0.52 |
|  |  | GS & TS | 2 | -0.001 | -0.09 | 0.08 |
| Ankle Dorsiflexor<br>(Weight Acceptance) | -0.02(0.01) | GS | 1 | -0.006 | -0.03 | 0.02 |
|  |  | GS & TS | 1 | -0.01 | -0.07 | 0.06 |
| Ankle Plantarflexor<br>(Controlled Lowering) | 1.16(0.20) | GS | 6 | 0.07* | -0.17 | 0.33 |
|  |  | GS & TS | 6 | 0.006 | -0.05 | 0.06 |

Slope coefficients (B) are reported for stair ascent gait speed (GS) and T-score (TS). Shaded areas indicate statistically significant findings, where the point estimate of the regression slope (B) was significantly different from zero at the following thresholds: $P \leq 0.05$ (\*), $**P \leq 0.01$ (\*\*), and $P \leq 0.001$ (\*\*\*).

### Joint moments

In the first regression model, stair descent speed significantly explained a portion of the variance in several peak joint moments (R² = 6–15%, P ≤ 0.05), including the hip abductor and knee extensor moments during weight acceptance, the hip extensor moment during the leg pull-through phase, and the hip flexor and ankle plantarflexor moments during the controlled lowering phase (Table 4, Figure 3). Including T-score in the second regression model did not increase the explained variance for any of the joint moment variables.

**Figure 3.**
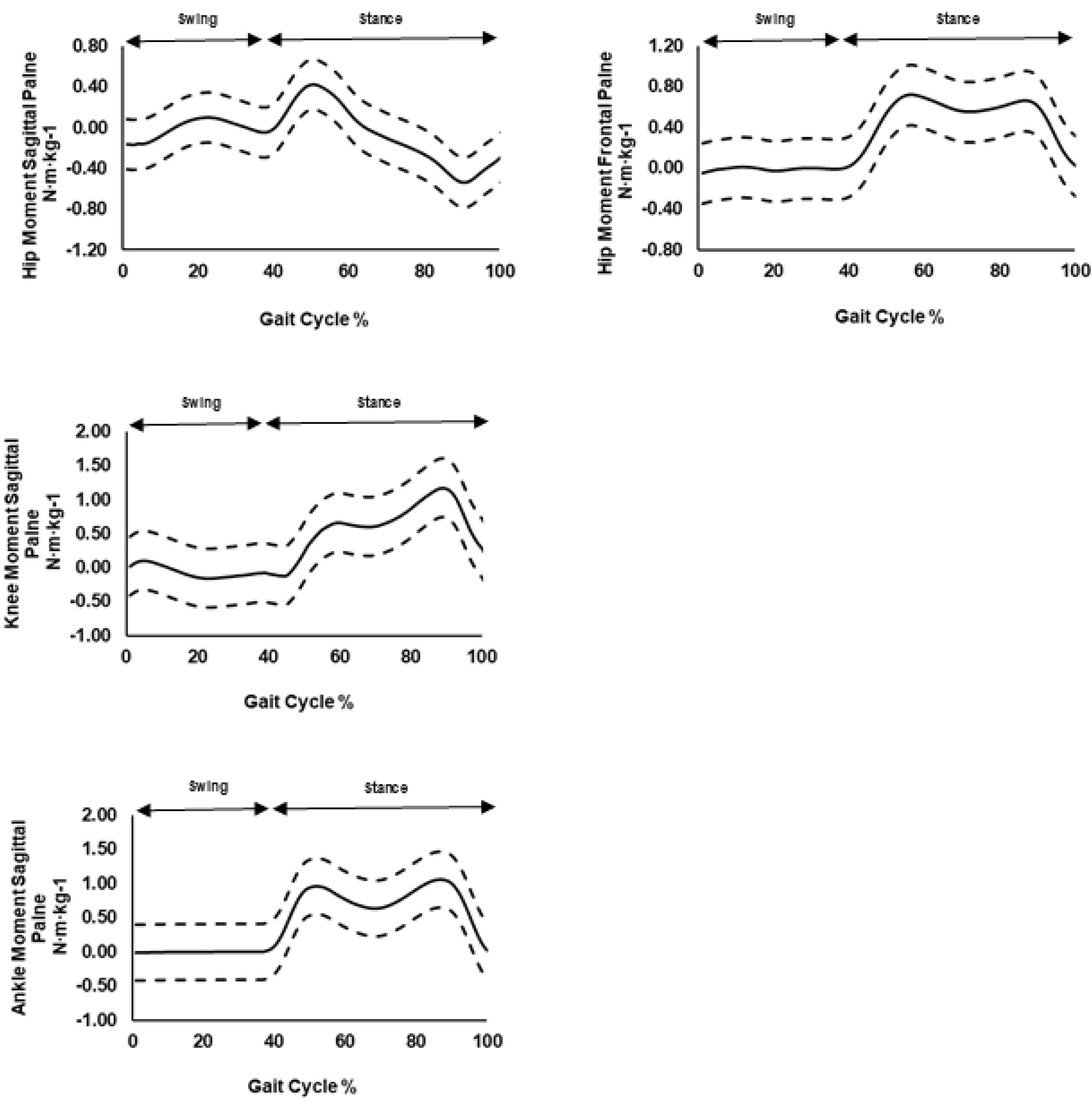
Ensemble averages (solid lines) ±1 standard deviation (dashed lines) of lower limb internal joint moments profiles during stair descent, normalised to body weight. Positive values represent hip abductor, hip and knee extensor, and ankle plantarflexor moments.

### Joint powers

Stair descent speed was also a strong predictor of variance in key joint power bursts (P ≤ 0.01), explaining 21% of the variance in K1 (knee extensor power absorption during weight acceptance), 19% in K2 (knee extensor power generation during forward continuance), and 49% in K5 (knee flexor power absorption during leg pull-through/foot placement). At the ankle, speed accounted for 13% of the variance in A1 (ankle plantarflexor power absorption during weight acceptance) and 23% in A3 (ankle plantarflexor power generation during controlled lowering) (Table 5, Figure 4). The addition of T-score in the second regression model did not significantly improve the explained variance for any joint power variables.

**Figure 4.**
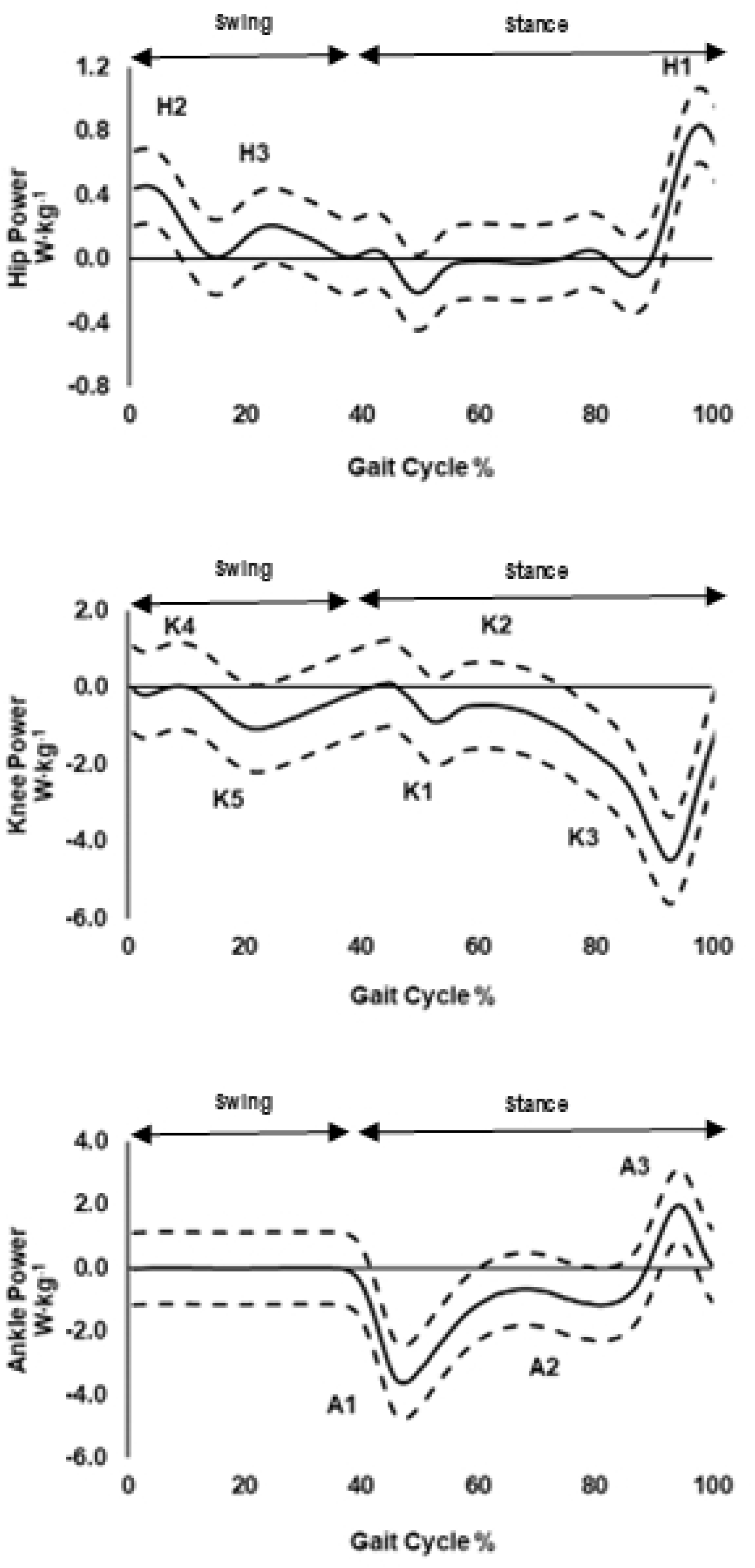
Ensemble averages (solid lines) ±1 standard deviation (dashed lines) of lower limb joint power profiles during stair descent. Distinct power bursts are labelled as H1–H3 (hip), K1–K4 (knee), and A1–A3 (ankle), following the convention established by McFadyen and Winter (1988). Positive joint power values indicate concentric (power-generating) muscle actions, while negative values represent eccentric (power-absorbing) actions.

**Table 5.**
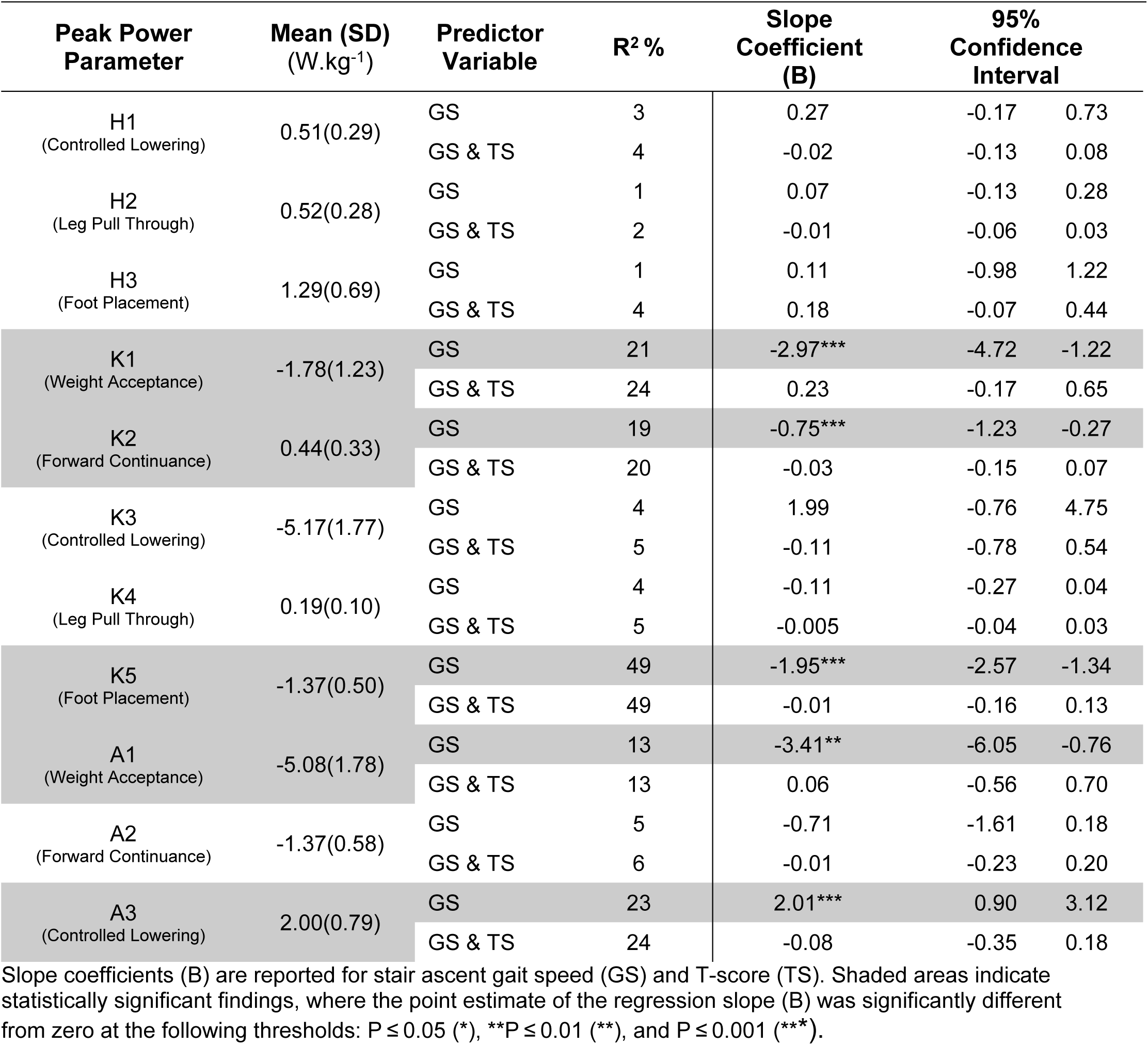
Explained variance (R²) and slope coefficients (B) for peak joint power during stair descent.

## Discussion

The purpose of this study was to quantify the extent to which stair descent speed and femoral neck T-score explained variability in lower limb gait parameters during stair descent in postmenopausal women across a wide range of bone mineral densities (from healthy to osteoporotic). The findings support our hypothesis, highlighting stair descent speed as a major explanatory factor, while T-score also contributed to the variance in specific kinematic and kinetic parameters. These results underscore the importance of accounting for both gait speed and BMD status when assessing older women during stair descent studies. Reporting mean descent speed is essential for meaningful between-study comparisons and for identifying early functional decline in clinical populations.

It should be acknowledged that the mean self-selected stair descent speed of our participants was 0.80 ± 0.21 m·s⁻¹, which is notably faster than the average speed reported for age-matched adults (0.65 m·s⁻¹) [33]. Our participants mean cadence during stair descent (112 ± 20 steps·min⁻¹), closely aligns with values typically reported for younger adults (110 ± 10 steps·min⁻¹). Despite some participants having reduced bone mineral density, their gait performance suggests a relatively high level of musculoskeletal function. This may be partially attributed to their self-reported engagement in physical activity, averaging five days per week (**Error! Reference source not found.**).

### Joint kinematics

Stair descent gait speed explained the variance in many kinematic parameters (Table 3). The inclusion of femoral neck T-score into the regression model significantly improved the explanatory power for anterior pelvic tilt, hip adduction, hip extension, knee flexion (P ≤ 0.01) and ankle dorsiflexion (P ≤ 0.001) during the controlled lowering and leg pull-through phases (Table 3). These findings suggest that reduced BMD was associated with altered movement strategies during stair descent. In particular, increased frontal plane motion at the hip and pelvis, stabilised primarily by the gluteus medius and other abductors, is critical for maintaining balance during stair descent, where greater stability demands are imposed compared to level walking [13,34]. During controlled lowering, the supporting limb must eccentrically control the descent of body mass, placing high demand on the knee extensors and requiring coordinated movement at the hip to preserve frontal plane stability. Although anterior pelvic tilt contributes to this control, excessive tilt displaces the centre of mass forward, increasing the risk of forward falls. These results underline the importance of hip, knee and pelvic muscle strength for safe stair descent, especially in older women with low BMD, where reduced muscular control could contribute to instability and fall-related injury. Moreover, during this phase, the knee extensors undergo substantial power absorption (i.e., eccentric contraction) to control whole body descent. These eccentric forces create mechanical loading on the femur, especially at the proximal region, which has been suggested to play an important role in preserving or enhancing BMD [3]. As such, regular stair descent in active older adults may improve dynamic stability and support musculoskeletal health by maintaining knee extensor strength and promoting osteogenic loading of the femur.

Our participants demonstrated approximately 4° more ankle dorsiflexion during controlled lowering compared to age-matched older adults [2,14]. Increasing ankle dorsiflexion during this phase of stair descent enhances energy generation by the gastrocnemius and soleus to stabilise the ankle [35,36]. Greater dorsiflexion allows the foot to maintain a flat contact with the step for longer, thereby increasing support and stability at a critical phase of the task [3]. This could reflect a deliberate adaptation to enhance safety by prioritising stability over propulsion.

### Ground reaction forces

The inclusion of femoral neck T-score as a predictor in the regression model significantly increased the explained variance in the mid-stance vertical GRF during forward continuance and the second vertical GRF peak during controlled lowering (both P ≤ 0.01) (Table 4). This suggests that T-score (BMD status) may influence how force is absorbed and managed during stair descent, particularly in phases requiring eccentric control of the lower limb.

The mid-stance GRF during forward continuance occurs when the body is transitioning between peak forces and is influenced by dynamic balance and shock absorption [37]. The significant contribution of T-score to explain the variance in this measure suggests that individuals with lower BMD adapt their gait mechanics, potentially through mechanisms such as altered joint stiffness [15] or muscle recruitment patterns to maintain postural stability and reduce impact loading during this transition phase. While these specific mechanisms were not directly measured, they warrant further investigation.

Specifically, the second GRF peak during controlled lowering reflects the force transmitted through the limb as the body is lowered onto the step below, placing high mechanical demands on the knee and ankle [38]. Lower T-scores, which indicate reduced BMD, may lead individuals to adopt more conservative (i.e., “safer”) loading strategies, potentially to reduce falls risk and the risk of injury. This could manifest as altered timing or reduced magnitude of GRF peaks.

### Joint moments

Stair descent speed explained the variance in most joint moments (Table 4 Table 3), including the hip extensor (P ≤ 0.001), hip flexor (P ≤ 0.05), knee extensor (P ≤ 0.01), and ankle plantarflexor (P ≤ 0.05) moments. This finding is consistent with our previous studies and the existing literature demonstrating that gait speed is a primary determinant of joint loading during dynamic tasks such as level walking and stair negotiation [9,24,31,32]. Notably, hip extensor and knee extensor moments showed the strongest associations (P ≤ 0.001 and P ≤ 0.01, respectively), underscoring the biomechanical demands placed on the knee extensors especially during the eccentric phases of stair descent.

In stair descent, particularly during the controlled lowering phase, the stance limb undergoes significant eccentric contraction during single limb support to modulate body mass as the individual steps downward. For our post-menopausal women, the hip flexors and knee extensors played an important role in resisting gravitational acceleration and ensuring controlled descent. The significant association between stair descent speed, hip flexor and ankle plantarflexor moments further illustrates the integrated muscular coordination required across the lower limb. While the hip flexors contribute to lifting and advancing the swing limb, the plantarflexors (e.g., gastrocnemius and soleus) help stabilise the ankle and control forward progression of the centre of mass [28]. At higher speeds, these muscles must act more forcefully or rapidly to preserve smooth motion and reduce the risk of missteps.

### Joint powers

The knee and ankle exhibited the most consistent and pronounced relationship with stair descent speed. Specifically, faster descent speeds were associated with increased power absorption during K1 (weight acceptance) and K5 (leg pull through/foot placement), and increased power generation during K2 burst (forward continuance) (Table 5) (Figure 4). These results reflect the complex role of the knee joint during stair descent, where eccentric contraction of the knee extensors is required to control body mass immediately after initial contact, and concentric action contributes to lifting the limb into swing later in the cycle. The K2 burst is normally indicative of knee extensor power generation in forward continuance, but our data were not consistent with McFadyen and Winter (1988) and indicated variability in the dataset. This was likely related to the broad range of our participants’ stair descent speeds and T-scores.

Importantly, these mechanical demands are heightened with faster speeds. The increased demands on the lower limb joints necessitate sufficient muscular strength and neuromuscular coordination. For postmenopausal women, particularly those with reduced BMD, the ability to meet these demands may be compromised, increasing the risk of instability or falls.

Similarly, stair descent speed significantly explained variance in A1 (weight acceptance) and A3 (controlled lowering) ankle power bursts (Table 5) (Figure 4). During weight acceptance, the ankle plantarflexors absorb power while during controlled lowering they generate power to relieve the extreme ankle dorsiflexion position and help push the swing leg through [28]. Higher ankle power requirements with increased speed suggest that ankle strength and control are important components of stair descent performance, and deficits here may lead to compensations further up the kinetic chain (e.g., at the knee, hip or pelvis).

The mechanical loading associated with increased joint power, particularly at the knee, may have positive implications for bone health. Eccentric muscle contractions apply tensile forces on the muscle-tendon complex which results in compressive loading to the bone, potentially stimulating osteogenic adaptation, which is crucial in populations with the propensity to develop osteopenia [39].

### Limitations

Several limitations of this study should be acknowledged. First, the participants appeared to be relatively high functioning for their age group, which may limit the generalisability of the findings to the broader population of older postmenopausal women, particularly those with lower physical function. Second, due to the cross-sectional design, causality cannot be established. Although associations between BMD and gait parameters were identified, longitudinal studies are needed to determine whether changes in gait mechanics result from low BMD or contribute to it.

While our study was sufficiently powered to detect moderate effect sizes (f² = 0.18) with the current sample size of 45 participants and two predictor variables, it may have been underpowered to detect small effects, particularly across the multiple dependent variables included in the analysis. Future studies should be adequately powered to improve sensitivity and allow for the detection of smaller but potentially meaningful effects, especially when examining complex or subtle relationships across multiple outcomes.

Another limitation is the reliance on self-reported number of days of being physically active, which is susceptible to recall bias and overestimation. Future studies should incorporate objective measures such as accelerometers or step counters to accurately assess daily activity levels, including stair use (e.g., number of floors ascended/descended and measured using wearables). Understanding the relationship between habitual loading activities and stair gait biomechanics could offer further insight into musculoskeletal adaptation in older women.

Finally, while self-selected gait speed reflects real-world behaviour, it also introduces inter-individual variability. Employing a protocol where participants walk at standardised speeds (e.g., slow, preferred, and fast) may help isolate the specific effects of speed on stair gait mechanics and improve the comparability of results across studies.

## Conclusions

This study highlights the complex interplay between gait speed, BMD, and lower limb biomechanics during stair descent in postmenopausal women. Our findings demonstrate that both stair descent speed and femoral neck T-score significantly influence joint kinematics, kinetics, and ground reaction forces, particularly during the critical controlled lowering phase. Reduced BMD was associated with altered movement strategies, including increased frontal plane motion at the hip and pelvis, which may serve to maintain balance but could also elevate the risk of instability and falls. The substantial eccentric muscle contractions, especially during the forward continuance and controlled lowering phases, generate mechanical loading on the lower limb bones, which can contribute to maintaining bone health.

This study highlights stair descent as not only a challenging functional task but also a potentially osteogenic activity when performed regularly and safely. Future research should further explore targeted interventions that enhance eccentric muscle capacity and joint stability to promote safe stair negotiation and preserve bone health in at-risk populations.

## Notes

### Competing Interest Statement

The authors have declared no competing interest.

